# Reconstitution of phospho-regulated mitotic chromatid assembly and disassembly

**DOI:** 10.1101/2025.09.08.674995

**Authors:** Keishi Shintomi, Shoji Tane, Yuki Masahara-Negishi, Tatsuya Hirano

**Affiliations:** Chromosome Dynamics Laboratory, RIKEN Pioneering Research Institute, Wako, Saitama, Japan

**Keywords:** Chromosome/Condensin/Mitosis/ Phosphorylation/Reconstitution

## Abstract

Exactly how cell cycle regulators control a sequential series of mitotic events is not fully understood. Here we report reconstitution assays that recapitulate the assembly and disassembly of mitotic chromatids in vitro using a minimal set of recombinant proteins, including condensin I. By incorporating cyclin B-Cdk1 and PP2A-B55, this system enables us to dissect the phospho-regulation of condensin I in these processes. We provide evidence that the terminal intrinsically disordered regions (tIDRs) of the non-SMC subunits suppress condensin I activity, and that this self-suppression is relieved by Cdk1 phosphorylation. Importantly, full activation of condensin I requires the phosphorylation of a conserved residue located in the central region of the kleisin subunit CAP-H. Conversely, PP2A-B55 induces dissociation of condensin I from reconstituted chromatids, leading to their disassembly. Complementary analyses using *Xenopus* egg extracts reveal that the tIDRs and the kleisin central region are phosphorylated and dephosphorylated with distinct kinetics during mitotic entry and exit. Together, these findings define an intricate regulatory network that coordinates chromatid assembly and disassembly with mitotic progression.

## Introduction

Chromosomes undergo large-scale structural transformations during mitosis. During mitotic entry, the amorphous mass of interphase chromatin is progressively converted into a discrete set of mitotic chromosomes, each composed of two rod-shaped sister chromatids, which ensures their accurate segregation into daughter cells. During mitotic exit, these rod-shaped chromatids are remodeled back into interphase chromatin as the cell nucleus reassembles, thereby restoring a chromatin state compatible with transcription and DNA replication in the subsequent cell cycle. These two transitions, referred to as mitotic chromatid assembly and disassembly, are thought to be under the strict control of cell cycle regulators. Elucidating their intricate regulatory mechanisms remains a major challenge in cell biology (Paulson *et al*, 2021; Vagnarelli, 2021; Dekker & Dekker, 2022; Hirano, 2025).

Extensive studies over the past three decades have established the consensus that a class of protein complexes, known as condensins I and II, plays a central role in the assembly of mitotic chromatids (Hirano, 2016; Hoencamp & Rowland, 2023). These two complexes share the same pair of SMC (structural maintenance of chromosomes) ATPase subunits but contain discrete sets of non-SMC subunits, and together form large protein assemblies of ∼650 kDa (Hirano & Mitchison, 1994; Saitoh *et al*, 1994; Hirano *et al*, 1997; Ono *et al*, 2003; Yeong *et al*, 2003). Recent advances in structural biology, single-molecule assays, and polymer physics-based modeling have begun to reveal how condensins harness the energy of ATP hydrolysis to drive dynamic conformational changes and reconfigure DNA structures, ultimately leading to mitotic chromatid assembly (Goloborodko *et al*, 2016; Ganji *et al*, 2018; Gibcus *et al*, 2018; Lee *et al*, 2020, 2022; Shaltiel *et al*, 2022).

With respect to the regulation of condensins, early studies revealed that condensin I is phosphorylated in a mitosis-specific manner (Hirano *et al*, 1997) and that this modification stimulates its ability to introduce positive DNA supercoils in vitro (Kimura *et al*, 1998; Takemoto *et al*, 2004; St-Pierre *et al*, 2009). Equally important, a mitotic chromatid reconstitution assay using a minimal set of purified factors demonstrated that phosphorylation by cyclin B-Cdk1 is sufficient to activate condensin I for chromatid assembly (Shintomi *et al*, 2015). Moreover, genetic studies have provided evidence that Cdk1 phosphorylation of the N-terminal tail of Smc4 regulates the nuclear import of condensin during mitosis in fission yeast (Sutani *et al*, 1999) and modulates the turnover of condensin on mitotic chromosomes in budding yeast (Robellet *et al*, 2015; Thadani *et al*, 2018). Despite these advances, the precise phosphorylation targets and the mechanistic aspects of condensin I activation remain poorly understood.

Far less is known about the regulation of chromatid disassembly during mitotic exit (Vagnarelli, 2021). Most mitotic phosphoproteins are rapidly dephosphorylated at the metaphase-anaphase transition, concomitant with cyclin B-Cdk1 inactivation. A major driver of this transition is protein phosphatase 2A complexed with its B55 regulatory subunit (PP2A-B55), which antagonizes cyclin B-Cdk1 activity by broadly dephosphorylating Cdk1 substrates to promote mitotic exit (Mochida *et al*, 2009; Godfrey *et al*, 2017; Rata *et al*, 2018; Kamenz *et al*, 2021; Chia *et al*, 2024). Although condensins are most likely dephosphorylated during mitotic exit, the responsible phosphatase(s) and their role in mitotic chromatid disassembly remain largely unexplored, primarily due to the lack of suitable experimental systems.

In the current study, we established in vitro functional assays using a minimal set of recombinant proteins, including condensin I, cyclin B-Cdk1, and PP2A-B55. These assays faithfully recapitulate both mitotic chromatid assembly and disassembly, and further allow us to dissect phosphorylation and dephosphorylation events that regulate condensin I function with unprecedented clarity. Together with complementary analyses using *Xenopus* egg extracts, our results demonstrate that the terminal intrinsically disordered regions (tIDRs) of the non-SMC subunits and a conserved residue located in the central region of CAP-H make distinct contributions to the assembly and disassembly of mitotic chromatids.

## Results

### Preparation of the complete set of fully recombinant proteins required for mitotic chromatid reconstitution

Mitotic chromatid reconstitution with a minimal set of purified factors provides a powerful assay to investigate the phospho-regulation of condensin I under near-physiological conditions. Among the six protein components used in the original assay, five (core histones, three histone chaperones, and topoisomerase II) were recombinant, whereas the mitotically phosphorylated form of condensin I was purified from *Xenopus* egg mitotic extracts (Shintomi *et al*, 2015). Although condensin I purified from interphase extracts failed to support this reaction, pre-phosphorylation with cyclin B-Cdk1 purified from starfish oocytes converted it into an active form. In our long-term effort to replace native condensin I and native cyclin B-Cdk1 with recombinant counterparts, we recently developed a protocol for producing an engineered version of recombinant cyclin B-Cdk1 (M-CDK) that is capable of mitosis-specific multi-site phosphorylation of various substrates, including condensin I (Shintomi *et al*, 2024). In the present study, we sought to combine the recombinant kinase with recombinant condensin I to establish a mitotic chromatid reconstitution assay composed entirely of recombinant proteins. During purification of the *Xenopus laevis* condensin I complex composed of full-length subunits (hereafter referred to as condensin I-FL; Fig. 1A), we found that the CAP-D2 and -H subunits were non-specifically phosphorylated in host insect cells, and that most of these modifications could be removed by λ-phosphatase treatment between the first and second chromatography steps (Fig. EV1A-C). To minimize potential effects of host-derived phosphorylation, all recombinant condensin I complexes used in this study were pre-treated with λ-phosphatase, which was removed in the final purification step.

**Figure 1.**
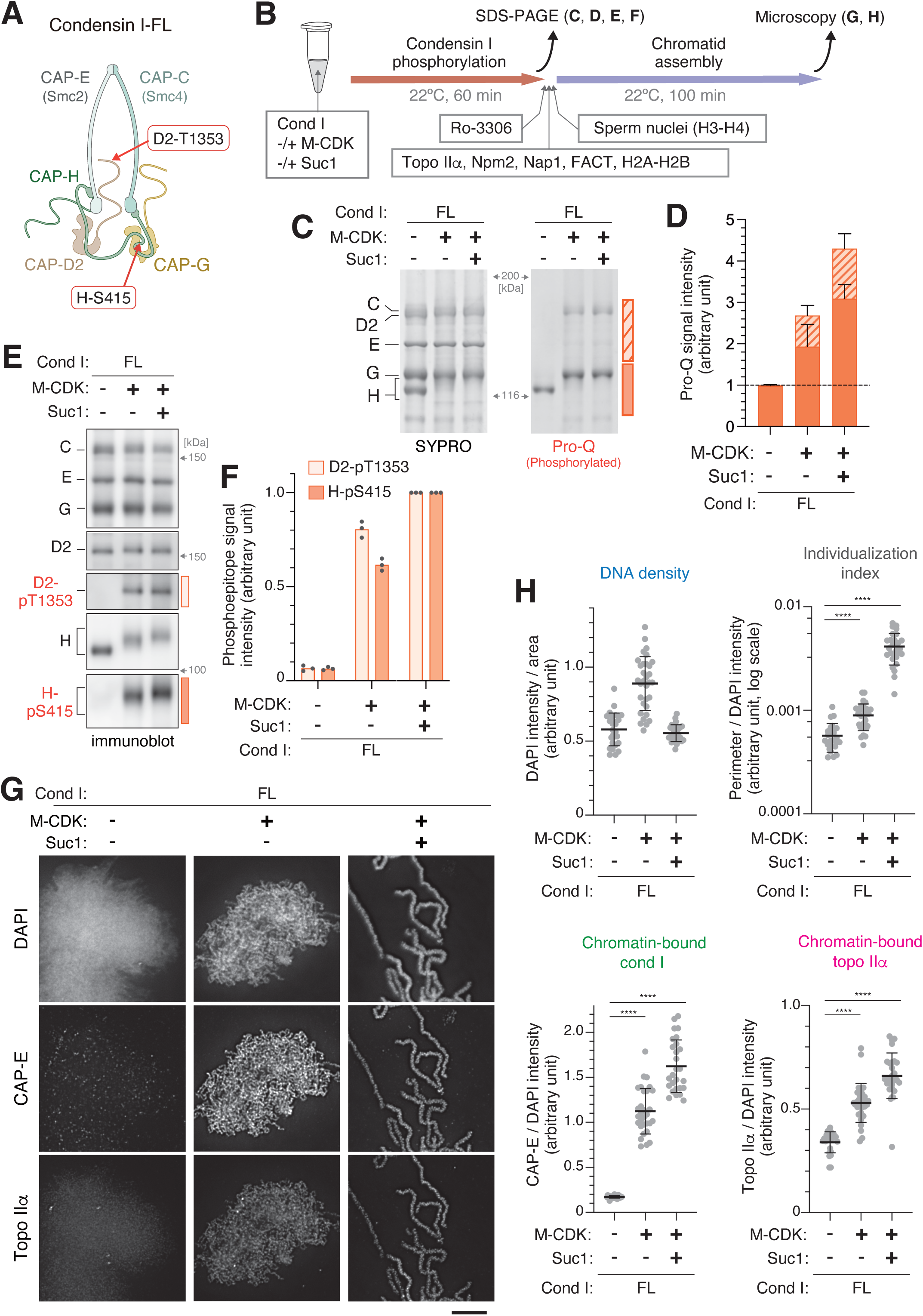
Cyclin B-Cdk1-Suc1 fully activates full-length condensin I in mitotic chromatid assembly assays. **A.** *Xenopus laevis* condensin I holocomplex composed of full-length subunits (condensin I-FL) and two Cdk1 phosphorylation sites analyzed in the current study (see also Fig EV1D). **B.** Scheme of condensin I phosphorylation followed by mitotic chromatid assembly. **C.** Condensin I-FL was either mock-treated (without kinase) or incubated with recombinant cyclin B-Cdk1 (M-CDK) in the absence or presence of Suc1 at 22°C for 60 min. The reaction mixtures were then analyzed by SDS-PAGE. The gel was first stained with Pro-Q Diamond to visualize phosphorylated proteins (right) and subsequently with SYPRO Ruby to visualize total proteins (left). **D.** The total intensities of Pro-Q Diamond signals in the upper and lower regions of the gel (indicated by striped and filled bars, respectively) were quantified. The values for reactions treated with M-CDK or M-CDK-Suc1 were normalized to those of the mock-treated control. **E.** The same set of reaction mixtures shown in (C) was analyzed by immunoblotting with antibodies against condensin I subunits and the phosphoepitopes pT1353 in CAP-D2 and pS415 in CAP-H. **F.** The intensities of the phosphoepitope signals were quantified. The values of the mock-treated and M-CDK-treated reactions were normalized to that of the M-CDK-Suc1-treated reaction. **G.** Reactions of mitotic chromatid assembly were set up as outlined in (B). Briefly, condensin I-FL was incubated with M-CDK and Suc1 in the indicated combinations, and phosphorylation was terminated by adding the Cdk1 inhibitor Ro-3306. The reactions were then mixed with a protein cocktail (topo IIα, histone chaperones [Npm2, Nap1, and FACT], and a truncated version of histone H2A.XF-H2B dimer), followed by the addition of *X. laevi*s sperm nuclei to initiate mitotic chromatid assembly. After incubation at 22°C for 100 min, the resulting chromatin structures were fixed, isolated, and labeled with antibodies against CAP-E (a condensin I subunit) and topo IIα. DNA was counterstained with DAPI. Scale bar 5 μm. **H.** Morphometric analyses of the chromatin structures shown in (G). DNA density (upper left) was calculated by dividing the integrated DAPI intensity by the area of each segment. The individualization index (upper right) was calculated by dividing the perimeter of each DAPI-positive segment by its integrated DAPI intensity. Chromatin-bound condensin I (lower left) and chromatin-bound topo IIα (lower right) were quantified by dividing the integrated signal intensity of CAP-E or topo IIα by that of DAPI in the same segment. Data information: In (D) and (F), data are presented as mean ± SEM from three independent experiments. In (H), data are presented as mean ± SD. **** *P*<0.0001 (two-tailed *t*-test with Welch’s correction).

### Cyclin B-Cdk1-Suc1 fully activates recombinant condensin I in chromatid reconstitution assays

As shown in our previous study (Shintomi *et al*, 2024), incubation of condensin I-FL with recombinant M-CDK alone resulted in a modest level of phosphorylation. The inclusion of Suc1, an adaptor protein that facilitates Cdk1 interaction with phospho-threonine-containing substrates (McGrath *et al*, 2013; preprint: Curran *et al*, 2025), markedly enhanced phosphorylation (Fig. 1B-F). We further found that M-CDK phosphorylated the CAP-C, -D2, and -H subunits of condensin I-FL, notably inducing a clear phosphorylation-dependent mobility shift of CAP-H (Fig. 1C). Quantification of the Pro-Q Diamond signals revealed that treatment with M-CDK alone increased the total phosphorylation level of condensin I-FL by ∼2.7-fold compared to the mock-treated control, whereas the addition of Suc1 led to a ∼4.3-fold increase (Fig. 1D). To evaluate site-specific phosphorylation of condensin I, we focused on two sites among the 18 Cdk consensus sites (SP and TP motifs) conserved between *X. laevis* and humans. D2-T1315, located in the C-terminal tail of CAP-D2 (Kimura *et al*, 1998), is representative of phosphorylation sites clustered in the terminal intrinsically disordered regions (tIDRs) of the non-SMC subunits, whereas H-S415, located in the central region of CAP-H (Kschonsak *et al*, 2017), represents the “core” of the condensin I complex (Figs. 1A and EV1D). Immunoblotting with the corresponding phospho-specific antibodies showed that both sites were phosphorylated by M-CDK, and that phosphorylation of these sites was further enhanced in the presence of Suc1 (Fig. 1E, F). Collectively, these assays enabled us to generate condensin I-FL complexes exhibiting distinct phosphorylation states, both quantitatively and qualitatively.

We next assessed the ability of these distinctly phosphorylated condensin I-FL preparations to assemble mitotic chromatids in the reconstitution assay. To eliminate potential contributions from phosphorylation of proteins other than condensin I, the Cdk1-specific inhibitor Ro-3306 was added to the reaction mixture together with the remaining components: topoisomerase IIα (topo IIα), core histones, three histone chaperones (Npm2, Nap1, and FACT), and sperm nuclei (Fig. 1B)(Shintomi & Hirano, 2021). After a 100-min incubation, chromatin was fixed, isolated, and subjected to immunofluorescent labeling using antibodies against CAP-E (a condensin I subunit) and topo IIα, along with DAPI counterstaining for DNA visualization. When condensin I-FL had not been treated with M-CDK, the sperm nuclei were decondensed and deformed by the actions of Npm2 and topo IIα, displaying an amorphous, cloud-like appearance with chromatin-bound condensin I below the detection limit (Fig.1G[left], H). This “null” phenotype was identical to that observed in reactions either lacking condensin I or using native condensin I prepared from interphase egg extracts (Shintomi *et al*, 2015). In contrast, moderate phosphorylation with M-CDK alone enabled condensin I-FL to associate with chromatin and form microscopically discernible fibrous structures with relatively high DNA densities. Under this condition, no individualized, rod-shaped chromatids were observed (Fig. 1G[middle], H). Strikingly, however, when condensin I-FL phosphorylated by M-CDK in the presence of Suc1 was used, its binding to chromatin was further enhanced, resulting in the formation of well-individualized, rod-shaped chromatids (Fig. 1G[right], H). It is noteworthy that, under the tested conditions, the amount of chromatin-bound topo IIα increased roughly in proportion to that of condensin I (Fig. 1G, H). Together, these results demonstrate that the ability of recombinant condensin I-FL to assemble mitotic chromatids in reconstitution assays is progressively enhanced through multisite, multisubunit phosphorylation mediated by M-CDK.

### Deletion of the tIDRs of the non-SMC subunits results in abortive or premature activation of condensin I

To investigate which subunits are critical for the M-CDK-mediated activation of condensin I, we next performed molecular dissection. From a structural point of view, the condensin I holocomplex consists of the core and the tIDRs extending from the non-SMC subunits. Among these, the C-terminal IDR of CAP-D2 (D2 C-tail) and the N-terminal IDR of CAP-H (H N-tail) harbor multiple consensus sites for Cdk phosphorylation (Fig. EV1D)(Kimura *et al*, 1998; Tane *et al*, 2022). To assess their roles in regulating condensin I function, we generated a truncation mutant lacking the three tIDRs of the non-SMC subunits, referred to as condensin I-ΔtIDRs (Figs. 2A and EV1D).

**Figure 2.**
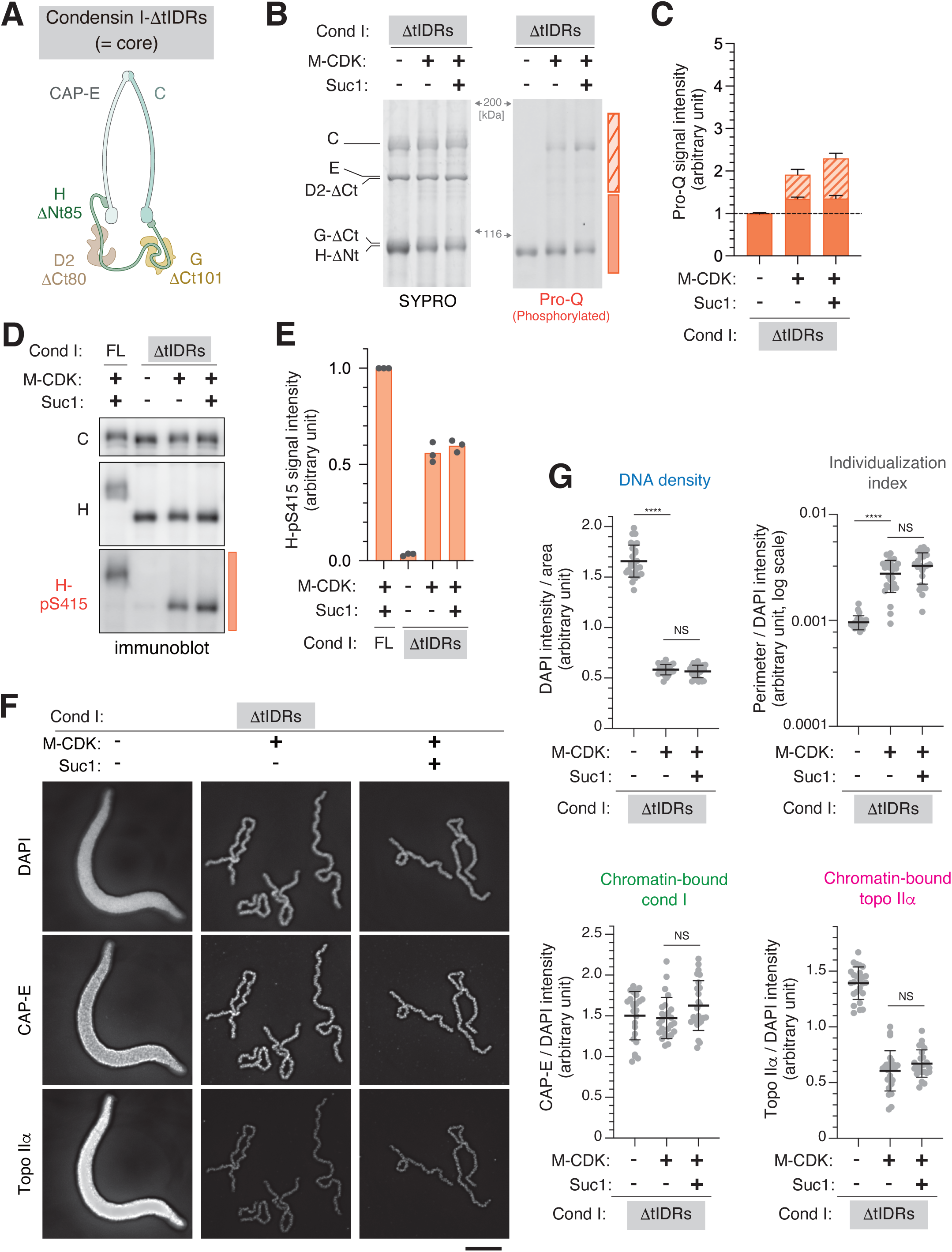
Deletion of the tIDRs of the non-SMC subunits results in abortive or premature activation of condensin I. **A.** *X. laevis* condensin I holocomplex lacking terminal intrinsically disordered regions (condensin I-ΔtIDRs), which we refer to as the condensin I “core” in this study. To prepare condensin I-ΔtIDRs, the N-terminal 85 amino acids of CAP-H, the C-terminal 80 amino acids of CAP-D2, and the C-terminal 101 amino acids of CAP-G were deleted (see also Fig. EV1D). **B-E**. Condensin I-ΔtIDRs was incubated with M-CDK and Suc1 in the indicated combinations. The reaction mixtures were analyzed as described in Fig. 1C-F. **F, G**. Condensin I-ΔtIDRs complexes characterized in (B-E) were subjected to mitotic chromatid assembly assays and analyzed as described in Fig. 1G, H. Scale bar 5 μm. Data information: In (C) and (E), data are presented as mean ± SEM from three independent experiments. In (G), data are presented as mean ± SD. **** *P*<0.0001, NS: not significant (two-tailed *t*-test with Welch’s correction).

We first examined M-CDK phosphorylation of condensin I-ΔtIDRs both quantitatively and qualitatively (Fig. 2B-E). Although condensin I-ΔtIDRs was phosphorylated upon incubation with M-CDK alone, the increase in the total phosphorylation level as judged by Pro-Q staining was lower than that of condensin I-FL (Fig. 2B, C). Moreover, the phosphorylation-dependent mobility shift of full-length CAP-H observed by SDS-PAGE (Fig. 1C, E) was hardly detectable in the truncated form of CAP-H (Fig. 2B, D). These observations suggest that a substantial fraction of Cdk1 target sites reside within the tIDRs of the non-SMC subunits. Furthermore, inclusion of Suc1 in the phosphorylation reaction did not lead to increased complex-wide phosphorylation or site-specific phosphorylation of H-S415 in condensin I-ΔtIDRs (Fig. 2B-E), in contrast to the pronounced enhancement observed in condensin I-FL (Fig. 1C-F).

We then tested the ability of these differentially treated condensin I-ΔtIDRs complexes to assemble mitotic chromatids in our reconstitution assay (Fig. 2F, G). We made two surprising observations in this experimental setup. First, unlike condensin I-FL without M-CDK treatment, untreated condensin I-ΔtIDRs did not display a null phenotype. Instead, it bound efficiently to chromatin and produced banana-shaped structures (Fig. 2F[left], G), rather than the amorphous, cloud-like structures observed with untreated condensin I-FL (Fig. 1G[left], H). Notably, a large amount of topo IIα was associated with these banana-shaped structures, despite their morphology resembling that observed in topo IIα-omitted reactions (Shintomi & Hirano, 2021). We speculate that, under this condition, condensin I-ΔtIDRs binds to chromatin in an abortive manner, thereby interfering with the proper action of topo IIα. Second, condensin I-ΔtIDRs treated with M-CDK alone acquired full activity to support the formation of individualized, rod-shaped chromatids (Fig. 2F[middle], G). The presence of Suc1 had little effect on either the morphology of the resulting chromatids or the amounts of chromosome-bound condensin I and topo IIα (Fig. 2F[right], G). These findings demonstrate that the tIDRs, when unphosphorylated, play an important role in preventing abortive chromatin binding by condensin I, and that their deletion lowers the threshold level of M-CDK phosphorylation required for full activation of condensin I. Moreover, the current results suggest that the core of condensin I contains key Cdk1 phosphorylation site(s), the modification of which not only suppresses the abortive action of condensin I-ΔtIDRs but also is required for its full activation.

### Phosphorylation of S415 in CAP-H is essential for full activation of condensin I

We next sought to investigate the importance of phosphorylation within the core of the condensin I complex. Among the potential phosphorylation sites, we focused on a Cdk consensus site located in the central region of CAP-H (S415 in *Xenopus*), which is conserved among most vertebrates (Fig. 3A). Our assay in fact confirmed that H-S415 is phosphorylated by M-CDK (Figs. 1E and 2D). To examine the functional contribution of this modification, we generated both full-length and ΔtIDRs versions of *Xenopus* condensin I carrying a phosphorylation-deficient mutation (S415A) in CAP-H (hereafter referred to as FL_H-S415A and ΔtIDRs_H-S415A, respectively). Quantitative and qualitative phosphorylation analyses (Fig. EV2A-D) reveal that overall phosphorylation levels in FL_H-S415A, both across the complex and at D2-T1353, were comparable to those of wild-type FL. In contrast, deletion of the tIDRs reduced the overall phosphorylation levels, even when combined with the S415A mutation, as observed in condensin I-ΔtIDRs (Fig. 2B-E).

**Figure 3.**
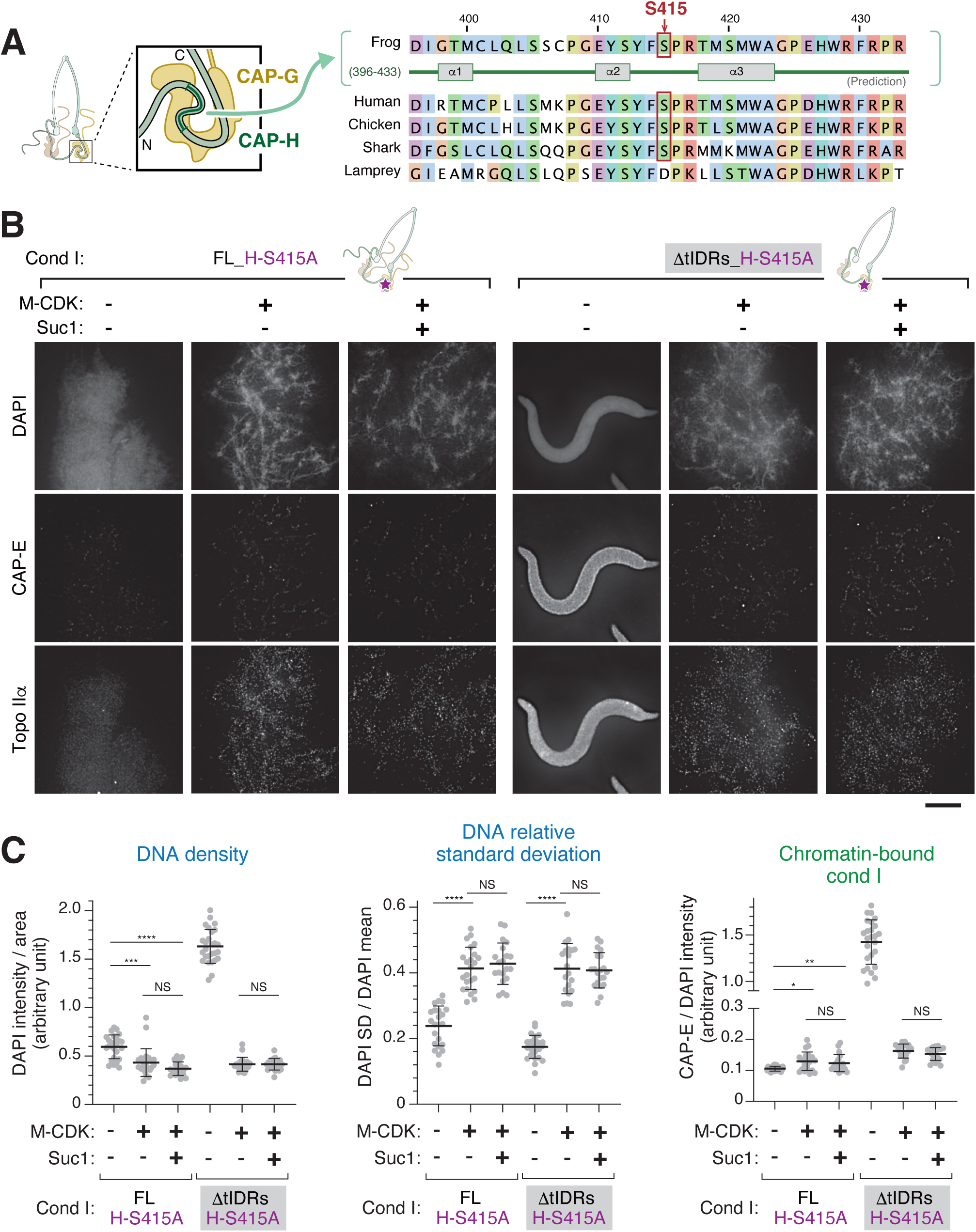
Phosphorylation of S415 in CAP-H is essential for full activation of condensin I. **A.** S415 in *X. laevis* CAP-H (frog) is located within the “safety belt” created by its interaction with CAP-G. An alignment of the corresponding regions of vertebrate CAP-H (human: *Homo sapiens*, chicken: *Gallus gallus*, shark: *Callorhinchus milii*, lamprey: *Petromyzon marinus*). See also Fig EV4. **B.** Condensin I-FL_H-S415A and condensin I-ΔtIDRs_H-S415 were mock-phosphorylated or phosphorylated by M-CDK in the absence or presence of Suc1 (Fig. EV3A-D) and then subjected to mitotic chromatid assembly assays. The resultant chromatin structures were analyzed as described in Figs. 1G and 2F (see also Fig. EV3E). **C.** DNA density (left) and chromatin-bound condensin I (right) were quantified as described in Fig. 1H. DNA relative standard deviation (center) was calculated by dividing the standard deviation of DAPI intensities within each segmented area by their mean value (see also Fig. EV3F). Scale bar, 5 μm. Data information: In (C), data are presented as mean ± SD. * *P*<0.05, ** *P*<0.01, *** *P*<0.001, **** *P*<0.0001, NS: not significant (two-tailed *t*-test with Welch’s correction).

In the absence of Cdk1 phosphorylation, condensin I-FL_H-S415A, like condensin I-FL, displayed a null phenotype in our reconstitution assay (Figs. 3B, C, and EV2E). However, when treated with M-CDK alone, condensin I-FL_H-S415A produced a previously unrecognized aberrant chromatin structure, characterized by discontinuous string-like structures surrounded by a diffuse chromatin cloud (Fig. 3B). Because it resembles a frosted, mist-like periphery, we refer to this unique morphological phenotype as “frost”. In the frost structures, the DAPI signal intensity was low and heterogeneous, as reflected by an increased relative standard deviation (Fig. 3C and EV2F). Only very low levels of the mutant complexes were detectable by immunofluorescence microscopy. Inclusion of Suc1 in the phosphorylation reaction had little effect on the resulting structures (Fig. 3B, C).

In the absence of Cdk1 phosphorylation, condensin I-ΔtIDRs_H-S415A, like condensin I-ΔtIDRs, aberrantly bound to chromatin and displayed the banana phenotype (Figs. 3B, C, and EV2E). When treated with M-CDK, regardless of whether Suc1 was included, the ΔtIDRs_H-S415A mutant generated frost structures virtually indistinguishable from those assembled with M-CDK-treated, condensin I-FL_H-S415A (Fig. 3B, C). These findings demonstrate that S415 in CAP-H is one of the most critical Cdk1 target sites required for full activation of condensin I. They also suggest that phosphorylation of site(s) in the core, in addition to H-S415, contributes to preventing the otherwise abortive chromatin binding of condensin I-ΔtIDRs.

### PP2A-B55 promotes condensin I dissociation from chromatids by dephosphorylating its tIDRs

Next, we sought to establish a mitotic chromatid disassembly assay by extending the chromatid reconstitution assay described above. We reasoned that PP2A-B55, a protein phosphatase known to dephosphorylate numerous Cdk1 targets, might act on Cdk1-phosphorylated condensin I and thereby inactivate the complex. To test this hypothesis, we expressed recombinant PP2A-B55 in insect cells and purified it to near homogeneity (Fig. 4A). We confirmed that this purified enzyme efficiently dephosphorylated an artificial substrate containing multiple TP motifs pre-phosphorylated by M-CDK, and that its activity was completely abolished by the potent PP2A inhibitor okadaic acid (Fig. EV3A).

**Figure 4.**
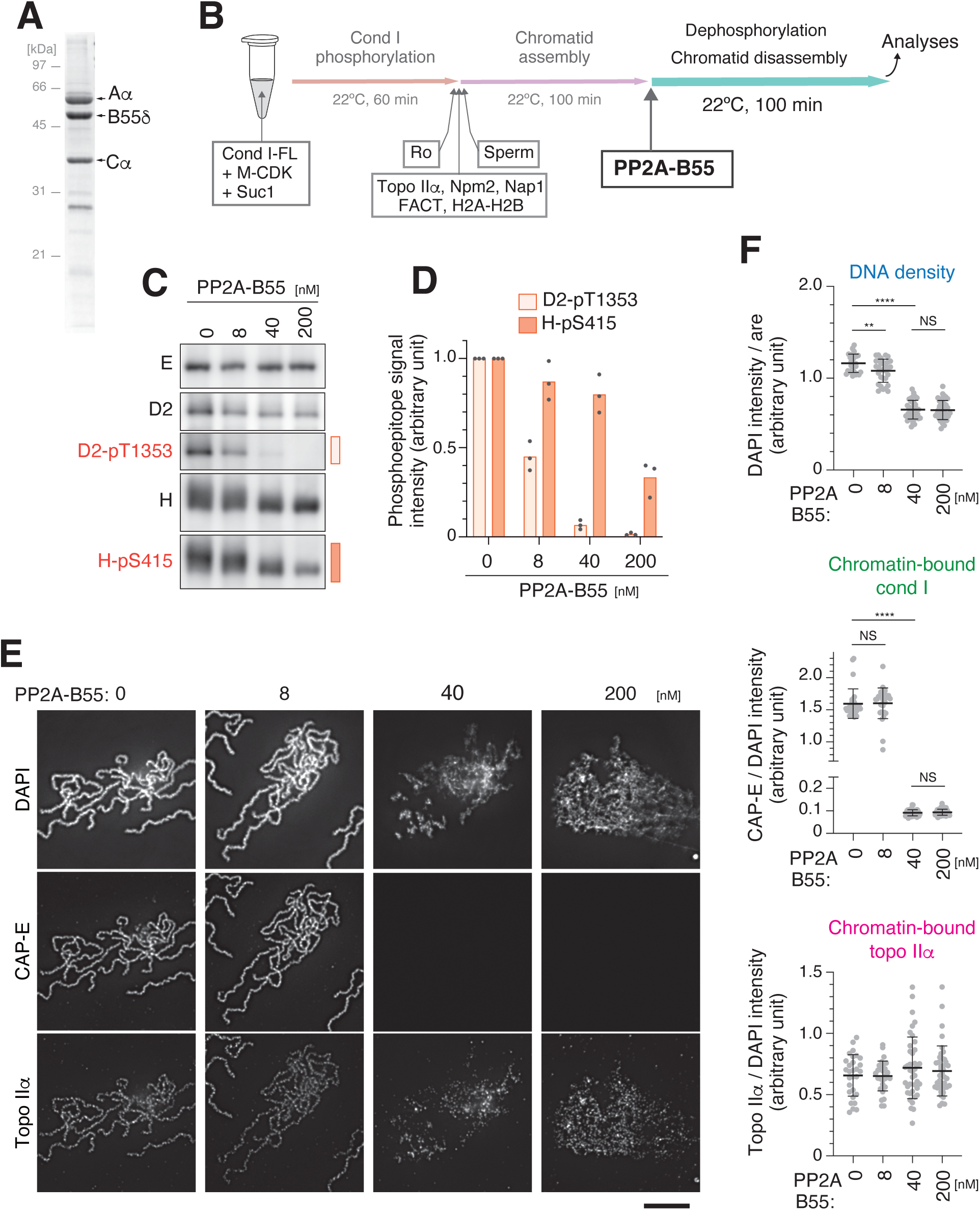
PP2A-B55 promotes condensin I dissociation from chromatids by dephosphorylating its tIDRs. **A.** The recombinant PP2A-B55 hetero-trimeric complex (comprising *Xenopus tropicalis* Aα, B55δ, and Cα subunits) was purified from insect cells and analyzed by SDS-PAGE followed by CBB staining. **B.** Scheme of mitotic chromatid disassembly assays. Briefly, mitotic chromatids were first assembled using condensin I-FL maximally phosphorylated by M-CDK in the presence of Suc1. The reaction mixture was then supplemented with increasing concentrations of PP2A-B55 and incubated for an additional 100 min. **C.** One hundred min after the addition of PP2A-B55, aliquots were taken and analyzed by immunoblotting with antibodies against condensin I subunits and the phosphoepitopes pT1353 in CAP-D2 and pS415 in CAP-H. **D.** Phosphoepitope signals in (C) were quantified and plotted, with values from PP2A-treated samples normalized to those from mock-treated controls. **E.** Chromatin structures in the PP2A-treated reactions were analyzed as described in Figs. 1G, 2F, and 3B. Scale bar, 5 μm. **F.** Morphometric parameters were quantified. See also Fig. EV3. Data information: In (F), data are presented as mean ± SD. ** *P*<0.01, **** *P*<0.0001, NS: not significant (two-tailed *t*-test with Welch’s correction).

We then added increasing amounts of PP2A-B55 to the reaction mixtures after completion of chromatid reconstitution (Fig. 4B). Titration experiments showed that T1353 in the tIDR of CAP-D2 was dephosphorylated more efficiently than S415 in CAP-H. At PP2A-B55 concentrations of 40 nM or higher, phosphorylation of D2-T1353 became almost undetectable (Fig. 4C, D). In parallel, microscopic observations revealed pronounced, dose-dependent deformation of originally rod-shaped chromatids (Fig. 4E). The deformation was accompanied by a reduction in DNA density and a marked loss of condensin I from chromatin. In contrast, the level of chromatin-bound topo IIα remained largely unchanged (Fig. 4F). The addition of okadaic acid together with PP2A-B55 blocked both condensin I dissociation from chromatin and chromatid disassembly, confirming the specificity of these reactions (Fig. EV3B). Thus, these results convincingly demonstrate that, in this experimental setup, addition of PP2A-B55 is sufficient to recapitulate mitotic chromatid disassembly in vitro, closely mirroring the process that occurs at the end of mitosis in cells.

### Sequential, site-specific phosphorylation and dephosphorylation of condensin I during mitotic entry and exit in *Xenopus* egg extracts

As described above, our reconstitution assays revealed that tIDR phosphorylation and core phosphorylation made distinct contributions to condensin I activation (Figs. 1-3). Moreover, dephosphorylation of D2-T1353 within the tIDRs was more closely associated with condensin I inactivation than that of H-S415 in the core (Fig. 4). To gain additional insight into how these functionally distinct phosphorylation and dephosphorylation events are temporally regulated in a physiological context, we turned to a *Xenopus* egg extract cell-free assay, which faithfully recapitulates cell cycle-dependent biochemical processes in a test tube (Murray, 1991; Shintomi & Hirano, 2017).

In the first set of assays, sperm nuclei were incubated in an interphase egg extract to allow nuclear assembly and DNA replication. Mitotic entry was then triggered by adding purified cyclin B protein. Immunofluorescence microscopy showed that nuclear envelope breakdown (NEBD) occurred synchronously around 30 min after cyclin B addition. Chromatin association of condensin I became apparent following NEBD (from 40 min onward), culminating in the formation of rod-shaped chromosomes by 100 min (Fig. 5A), as reported previously (Shintomi & Hirano, 2011). Time-course analysis of condensin I phosphorylation revealed that phosphorylation of D2-T1353 occurred relatively early (10-20 min), whereas phosphorylation of H-S415 became prominent only after 40 min (Fig. 5B). Quantitative analysis revealed a tight correlation between H-S415 phosphorylation and the accumulation of condensin I on chromatin (Fig. 5C), supporting the notion that mitotic activation of condensin I is achieved in a stepwise manner through sequential phosphorylation of the tIDRs followed by H-S415.

**Figure 5.**
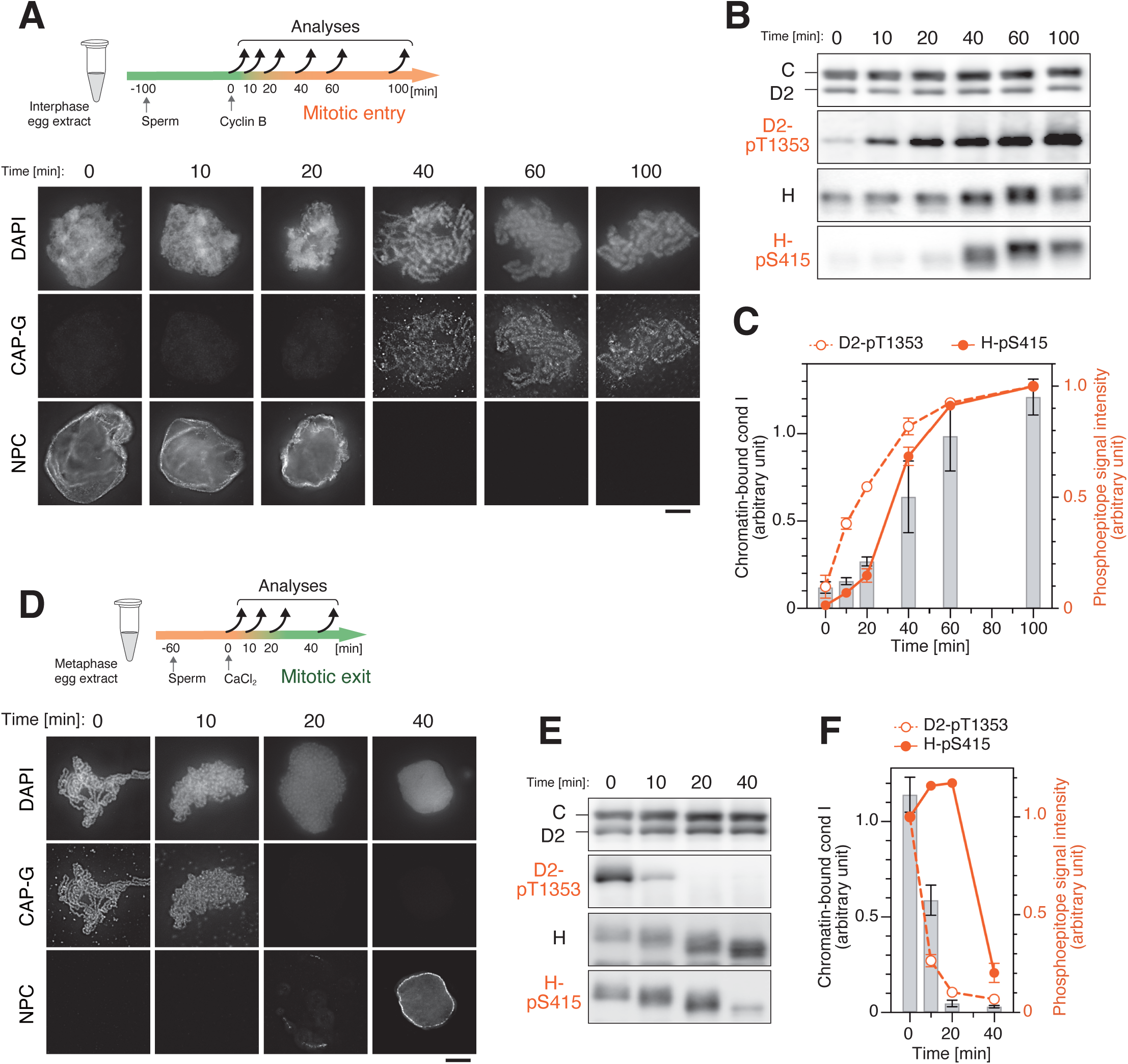
Sequential, site-specific phosphorylation and dephosphorylation of condensin I during mitotic entry and exit in *Xenopus* egg extracts. **A.** Sperm nuclei were first incubated with interphase egg extracts to allow nuclear assembly and DNA replication, and then cyclin B was added to trigger mitotic entry. At the indicated time points after cyclin B addition, nuclei and chromosomes were fixed and labeled with antibodies against CAP-G (a condensin I subunit) and components of the nuclear pore complex (NPC). **B.** In the same reaction mixtures as analyzed in (A), aliquots of the extract were taken at the indicated time points and analyzed by immunoblotting with the indicated antibodies. **C.** Chromatin-bound condensin I and phosphoepitope intensities were quantified and plotted. **D-F.** Sperm nuclei were first incubated with metaphase-arrested egg extracts to allow mitotic chromatid assembly for 60 min, and then CaCl_2_ was added to trigger the cell cycle transition into interphase. The samples were analyzed by immunofluorescence microscopy and immunoblotting as described in (A)-(C).

In the second set of assays, we examined how condensin I is inactivated during mitotic exit. To this end, sperm nuclei were first incubated in a metaphase-arrested extract, allowing the assembly of single chromatids with condensin I concentrated along their axes. Mitotic exit was then triggered by the addition of Ca^2+^ ions. Immunofluorescence microscopy showed that chromatid disassembly occurred 10-20 min after Ca^2+^ addition, concomitant with condensin I dissociation from chromatin, and was followed by nuclear envelope reformation at 40 min (Fig. 5D). We observed a rapid dephosphorylation of D2-T1353 at 10-20 min, whereas dephosphorylation of H-S415 occurred more slowly, at 20-40 min (Fig. 5E). Notably, the timing of condensin I dissociation from chromatin closely paralleled that of D2-T1353 dephosphorylation (Fig. 5F). These results are consistent with the reconstitution assays using defined factors, in which PP2A-B55-mediated dephosphorylation of D2-T1353 is coupled to condensin I inactivation.

## Discussion

In this study, we established in vitro reconstitution assays that recapitulate the assembly and disassembly of mitotic chromatids using a minimal set of fully recombinant proteins. These assays allow us to dissect how Cdk1-mediated phosphorylation and PP2A-mediated dephosphorylation activate and inactivate condensin I function, respectively, during mitotic entry and exit. Importantly, our results reveal that the ordered phosphorylation and dephosphorylation of the tIDRs and the core of condensin I regulate the complex through distinct mechanisms.

In the reconstitution assays reported here, we are now able to analyze a panel of wild-type and mutant forms (e.g., a tIDR deletion or point mutation) of condensin I, each subjected to distinct quantitative and qualitative levels of Cdk1 phosphorylation. The ability of these complexes to assemble mitotic chromatids in our reconstitution assays is summarized in Fig. 6A. One of the key findings is that the tIDRs of the non-SMC subunits, notably enriched in Cdk1 consensus sites (Fig. EV1D), play a critical role in condensin I regulation. The current results, together with our previous work (Shintomi *et al*, 2024), demonstrate that phosphorylation by M-CDK in the presence of Suc1 is essential for full activation of condensin I-FL (Figs. 1 and 6A): three TP motifs in the CAP-D2 tIDR (T1314, T1348, and T1353 in *X. laevis*) likely serve as Suc1 recognition sites once they are phosphorylated. Importantly, deletion of the tIDRs renders unphosphorylated condensin I out of control, thereby triggering abortive chromatin binding, and also attenuates the total phosphorylation level required for condensin I to assemble mitotic chromatids (Figs. 2 and 6A). We therefore suggest that the tIDRs of condensin I function as more than a simple self-suppressing module relieved by phosphorylation; rather, they constitute a more intricate regulatory element that prevents premature activation by sensing and responding to distinct phosphorylation states during cell cycle progression, in a manner reminiscent of for the Cdk inhibitor Sic1 (Nash *et al*, 2001). The current results obtained from our reconstitution assays complement and extend our previous findings in *Xenopus* egg extracts, in which deletion of the CAP-H N-terminal tail not only accelerated condensin I loading in mitotic extracts but also rendered it partially active even in interphase extracts (Tane *et al*, 2022). More recently, it has been reported that conserved short linear motifs (SLiMs) in the tIDRs of CAP-D2 and CAP-H interact with CAP-G to suppress the DNA-stimulated ATPase activity of condensin I (Cutts *et al*, 2025). The precise mechanisms by which each tIDR functions in our reconstitution assays remain to be clarified in future studies. It is worth noting that the tIDR-mediated self-suppression and release mechanisms discussed here also operate in the regulation of condensin II (Yoshida *et al*, 2022, 2024).

**Figure 6.**
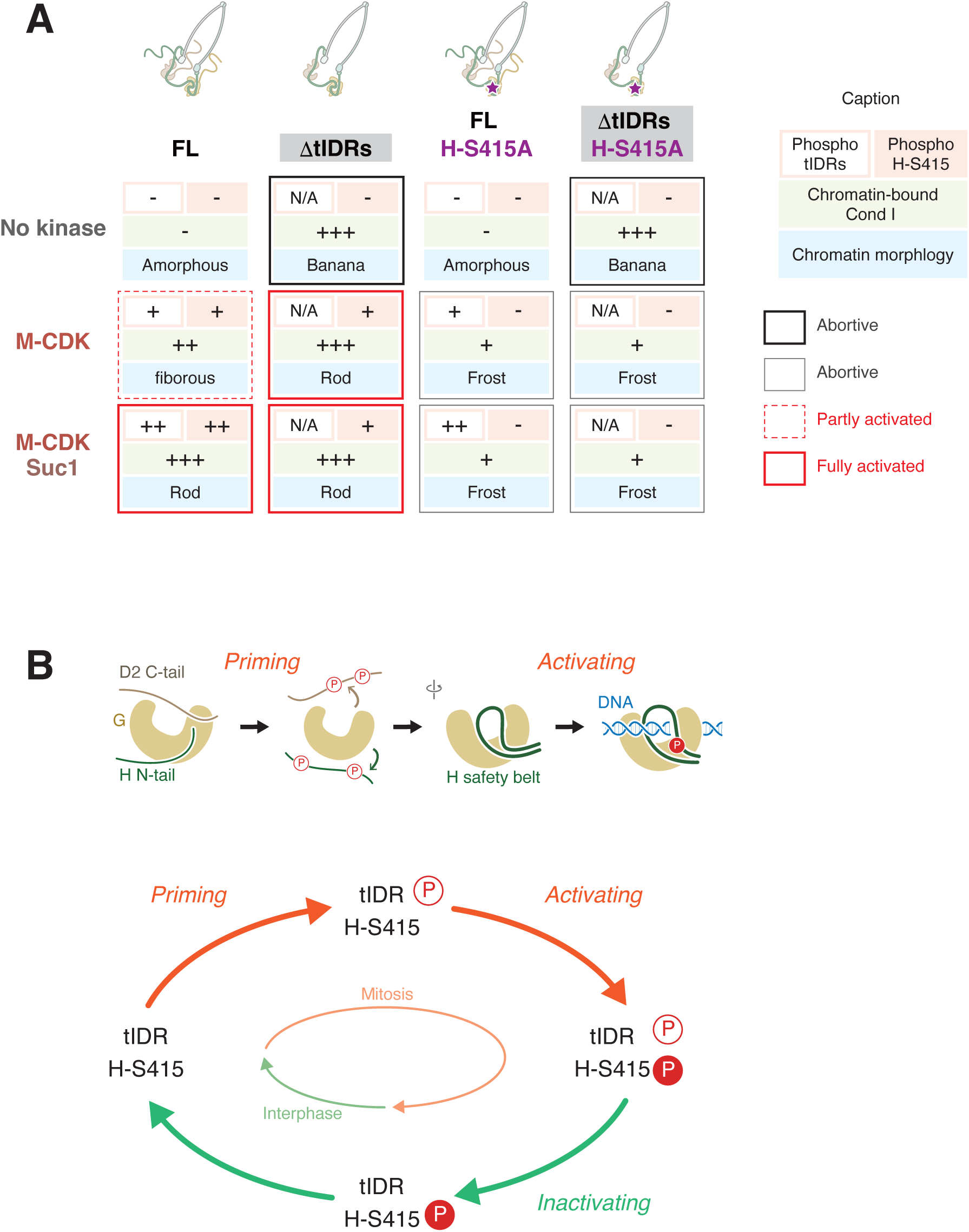
Phospho-regulation of condensin I. **A.** Summary of reconstitution assays for mitotic chromatid assembly shown in Figs. 1, 2, and 3. The updated reconstitution assays employing fully recombinant proteins enabled us to generate condensin I with distinct phosphorylation states. Without kinase treatment, the full-length (FL) complex exhibited a “null” phenotype (i.e., “amorphous” structures with no condensin I binding), whereas the ΔtIDRs complex bound to chromatin abortively, producing “banana”-shaped structures. A single phosphorylation-deficient mutation at S415 in CAP-H (H-S415A) severely impaired chromatin binding regardless of the presence of the tIDRs, leading to the “frost” phenotype. Thus, coordinated phosphorylation events in the tIDRs and at H-S415 are essential for full activation of condensin I and for the assembly of “rod”-shaped mitotic chromatids. **B.** Models for sequential, site-specific phospho-regulation of condensin I during the cell cycle. During interphase, the D2 C-tail and H N-tail (tIDRs) interact with CAP-G to suppress condensin I activity. At mitotic entry, cyclin B-Cdk1 first phosphorylates these tIDRs, relieving this suppression (“priming” phosphorylation). Subsequently, phosphorylation of S415 within the CAP-H safety-belt motif fine-tunes interactions between the anchor module and DNA, thereby promoting mitotic chromatid assembly (“activating” phosphorylation). At mitotic exit, PP2A-B55 first dephosphorylates the tIDRs, triggering condensin I dissociation from chromatin and mitotic chromatid disassembly. H-S415 is likely dephosphorylated later, returning condensin I to an interphase state. See Discussion for details.

Another important finding of this study is that phosphorylation of S415 in CAP-H is essential for full activation of condensin I, irrespective of the presence or absence of the tIDRs (Fig. 3; Fig. 6A). Our results are consistent with the previous observation that alanine substitution of the corresponding residue, S432 in human CAP-H compromised condensin I binding to mitotic chromosomes in HeLa cells (Kschonsak *et al*, 2017). Then how does the phosphorylation of the core segment of CAP-H regulate condensin I activity? Interestingly, S415 in *Xenopus* CAP-H (S432 in human CAP-H) is predicted within the so-called “safety belt”, which has been proposed to anchor one of the DNA segments involved in loop extrusion (Fig. EV4). A recent molecular dynamics simulation study suggests that tight closure of the safety belt may raise the energetic barrier for initial DNA entrapment achieved by the anchor module composed of CAP-H/Brn1 and CAP-G/Ycg1 (preprint: Chen *et al*, 2025). Because H-S415 lies close the “latch” of the safety belt, phosphorylation of this site is likely to modulate the belt’s conformation, either loosening or tightening, and thereby fine-tune condensin I’s interaction with dsDNA (Fig. 6B, upper). Finally, we note that the abortive chromatin binding of condensin I-ΔtIDRs can be reversed by M-CDK even in the presence of the H-S415A mutation (Figs. 3 and 6A), suggesting the existence of additional phosphorylation site(s) in the core that alter the functional properties of condensin I. Identifying these site(s) will further enrich our understanding of the highly intricate phospho-regulation of condensin I.

Compared with the mechanism of mitotic chromatid assembly, much less is known about chromatid disassembly during mitotic exit. To the best of our knowledge, the reconstitution assay reported here is the first to recapitulate this process using a defined set of fully recombinant proteins. Given previous reports implicating the Cdc48 ATPase (Thattikota *et al*, 2018) and RuvB-like ATPases (Magalska *et al*, 2014) in chromatid decondensation, we were somewhat surprised to find that treatment with PP2A-B55 alone was sufficient to trigger condensin I dissociation and subsequent chromatid disassembly in our reconstitution assay (Fig. 4). While we believe that condensin I is the major (and possibly sole) target of PP2A-B55 responsible for mitotic chromatid disassembly in this system, the current experimental setup does not exclude the possibility that concomitant dephosphorylation of other components in the protein cocktail contributes marginally to this process. Moreover, our results do not rule out the possible involvement of additional phosphatases (Vagnarelli *et al*, 2006; preprint: Zeisner *et al*, 2025).

As an approach complementary to the reconstitution assays, we employed *Xenopus* egg extracts, in which spatiotemporal progression of cell cycle events, including mitotic chromatid assembly and disassembly, can be recapitulated under more physiological conditions. The extract-based assay clearly showed that the two phosphorylation sites, D2-T1353 and H-S415, undergo phosphorylation and dephosphorylation with different kinetics. Notably, phosphorylation of D2-T1353 begins prior to nuclear envelope breakdown, whereas phosphorylation of H-S415 coincides with nuclear envelope breakdown and mitotic chromatid assembly (Fig. 5A-C). It is therefore conceivable that phosphorylation of the tIDRs places condensin I in a “primed” state, which is subsequently converted into a fully active state by phosphorylation of the core (Fig. 6B). Moreover, the priming phosphorylation is likely to occur in the cytoplasm, given that condensin I exclusion from the nucleus has recently been shown to be essential for preventing its premature activation in human cells (Eykelenboom *et al*, 2025). It is also noteworthy that dephosphorylation of D2-T1353 precedes that of H-S415 and coincides with mitotic chromatid disassembly (Figs. 5D-F and 6B). This reversible switch may function as a bistable module that integrates the opposing activities of Cdk1 and PP2A. Such a mechanism would be particularly advantageous at mitotic exit, when chromatid disassembly must be completed within a narrow time window.

Looking ahead, our reconstitution system provides a tractable platform to address several outstanding questions in the field. Its ability to recapitulate distinct phosphorylation states offers a unique opportunity to investigate how specific phosphorylation patterns influence condensin-mediated activities, including loop extrusion (Ganji *et al*, 2018), DNA-DNA capture (Tang *et al*, 2023), and functional interplay with topo II (Tsubota *et al*, 2025). Moreover, combining biochemical reconstitution with other cutting-edge techniques (e.g., cryo-electron microscopy and molecular dynamics simulations) could yield deeper insights into how phosphorylation and dephosphorylation rewire the structural and functional landscapes of condensins and related molecular machines.

## Methods

### Preparation of recombinant proteins

#### *Xenopus laevis* condensin I

Recombinant condensin I complexes were produced as reported previously (Shintomi *et al*, 2024) with some modifications. Briefly, cDNA fragments encoding the five subunits of condensin I (CAP-C, -D2, -E, -G, and -H) were codon-optimized for *Trichoplusia ni* and synthesized by a commercial provider (GeneArt, Thermo Fisher Scientific). Their full-length and truncated versions (CAP-D2_ΔCt80, CAP-G_ΔCt101, and CAP-H_ΔNt85, see Fig. EV1D) were amplified by PCR and cloned individually into the plasmid vector pLIB (Addgene, Catalog # 80610) or its lab-made derivatives (for fusing affinity tags). Site-directed mutagenesis to generate the CAP-H S415A substitution was performed using QuikChange Lightning Mutagenesis Kit (Agilent Technologies, 210518). DNA cassettes encoding 3xFLAG-tagged CAP-C and GST-tagged CAP-E (3xFLAG-tag and GST-tag are cleavable by TEV and 3C protease, respectively) were excised from the pLIB-based vectors and assembled into the pBIG1A vector (Addgene, 80611) using Gibson Assembly Cloning Kit (New England Biolabs, E5510). Corresponding DNA cassettes for full-length or mutant CAP-D2, CAP-G, and CAP-H were similarly integrated into the pBIG1B vector (Addgene, 80612). The resulting pBIG1-based vectors were introduced into the *Escherichia coli* strain DH10Bac (Thermo Fisher Scientific, 10361012) to generate recombinant bacmid DNAs via Tn7-mediated transposition. Baculoviruses were produced by transfecting Sf9 cells (Thermo Fisher Scientific, 11496015) with the bacmids and further amplified by an additional round of infection. High-Five cells (Thermo Fisher Scientific, B85502) were then co-infected with amplified viruses and grown in Express Five medium (Thermo Fisher Scientific, 10486025) at 27°C for 48 h. The cells were harvested and resuspended in buffer ST200 (20 mM Hepes-NaOH [pH 8.0], 200 mM NaCl, 10% glycerol) supplemented with EDTA-free Complete Protease Inhibitor Cocktail (Merck, 11873580001) and Benzonase nuclease (Merck, 71205), and then lysed by sonication. The lysate was clarified by centrifugation at 35,000 g for 15 min and applied to a Strep-Tactin XT column (IBA Lifesciences, 2-5031-010). The column was washed with buffer ST200, and bound proteins were eluted with 50 mM D-Biotin (Nacalai Tesque, 04822-91). The eluate was first dialyzed against buffer ST150 (20 mM Hepes-NaOH [pH 8.0], 150 mM NaCl, 10% glycerol) in the presence of lab-made 3C protease to cleave a dispensable GST-tag, and was then supplemented with lab-made λ-phosphatase and 1 mM MnCl_2_ to remove host-derived phosphorylation as much as possible (see Fig. EV1B-D). The resultant partially purified complex was mixed with anti-FLAG beads (Fujifilm Wako Pure Chemicals, 012-22384) and condensin I bound to the antibody was released by TEV-protease-mediated cleavage. Peak fractions were pooled, dialyzed against buffer KHG150/10 (20 mM Hepes-KOH [pH 7.7], 150 mM KCl, 10% glycerol) supplemented with 0.5 mM TCEP (Fujifilm Wako Pure Chemicals, 203-20153), and concentrated with an Amicon Ultra MWCO-100K device (Merck, UFC910024). The concentrated protein sample was dispensed into small aliquots, snap-frozen in liquid nitrogen, and stored at −80°C until use.

#### *Xenopus tropicalis* PP2A-B55

cDNA fragments for *X. tropicalis* PP2A-Aα (PPP2R1A), PP2A-B55δ (PPP2R2D), PP2A-Cα (PPP2CA) were codon-optimized for *Trichoplusia ni* and synthesized by a commercial provider (Eurofins Genomics). These fragments were sequentially cloned into plasmid vectors to generate a pBIG1-based vector harboring Twin-strep-tagged Aα and B55δ, and His6-tagged Cα. Bacmid DNA generation, baculovirus production, and protein expression were carried out as described for condensin I. The resultant PP2A-B55 complex was first purified with Ni^2+^-charged column and further purified on a Strep-Tactin column. Peak fractions were pooled, dialyzed against buffer KHG150/10 (20 mM Hepes-KOH [pH 7.7], 150 mM KCl, 10% glycerol) supplemented with 0.5 mM TCEP, and concentrated with an Amicon Ultra MWCO-30K device (Merck, UFC503024). The concentrated protein sample was dispensed into small aliquots, snap-frozen in liquid nitrogen, and stored at −80°C until use.

### Other proteins

M-CDK (an engineered version of cyclin B-Cdk1 complex from *X. tropicalis*), Suc1 (a Cks1 homolog in *Schizosaccharomyces pombe*), and MBP-XD2-C (a maltose-binding protein [MBP]-fused *X. laevis* CAP-D2 tIDR) were prepared as previously described (Shintomi *et al*, 2024). The concentrations of all recombinant proteins were determined by measuring absorbance at 280 nm.

### Phosphorylation of condensin I by M-CDK

A reaction mixture containing condensin I, M-CDK, and Suc1 in various combination was dialyzed against buffer PR (20 mM Hepes-KOH [pH 7.7], 80 mM KCl, 5 mM MgCl_2_) using an Xpress Micro Dialyzer MD100 (Scienova, 40078) at 4°C. Protein concentrations in the mixture were as follows: condensin I, 100 nM; M-CDK, 0 or 50 nM; Suc1, 0 or 500 nM. The dialysate was supplemented with 2 mM ATP and incubated at 22°C for 60 min to allow phosphorylation. Aliquots were taken and mixed with an SDS sample buffer. The samples were subjected to SDS-PAGE, followed by Pro-Q Diamond (Thermo Fisher Scientific, P33300) staining to visualize phosphorylated proteins and SYPRO Ruby (Thermo Fisher Scientific, P12000) to visualize total proteins. The same set of samples was also analyzed by immunoblotting to detect the phosphoepitopes pT1353 in CAP-D2 and pS415 in CAP-H.

After the phosphorylation assay, the resultant mixtures were also used in subsequent chromatid reconstitution assays as a source of phosphorylated condensin I. For this purpose, the mixtures were supplemented with the Cdk1 inhibitor Ro-3306 (ChemScene, CS-3790) at a concentration of 9 µM to prevent further phosphorylation of condensin I as well as other proteins during chromatid reconstitution.

### Reconstitution of mitotic chromatid assembly and disassembly in vitro

Mitotic chromatid reconstitution (mitotic chromatid assembly) assays were performed as previously described with some modifications. Briefly, the phosphorylation reaction mixture and other protein factors required for chromatid reconstitution (each dialyzed against buffer PR) were combined to achieve the following final concentrations: condensin I, 20 nM; topoisomerase IIα, 40 nM; Npm2, 60 µM; Nap1, 4.5 µM; dX-dB (a complex of histone H2A-H2B derivatives), 1.0 µM; and FACT, 360 nM. The reaction mixture was supplemented with 2 mM ATP, 10 mM phosphoenolpyruvate (Sigma, P0564), and 100 units/ml pyruvate kinase (Sigma, P9136). Chromatid reconstitution was initiated by adding demembranated *Xenopus* sperm nuclei (1,000 nuclei/µl) to the mixture (final volume: 20 µl), followed by incubation at 22 °C for 100 min.

Mitotic chromatid disassembly assays shown in Figs. 4 and EV4B were performed as follows: after completion of mitotic chromatid assembly as described above, the reaction mixtures were supplemented with PP2A-B55 at the indicated concentrations and incubated for an additional 100 min, either in the presence or absence of okadaic acid (1 µM). Finally, the reaction mixtures were fixed and subjected to immunofluorescence microscopy.

### Validation of recombinant PP2A-B55 (Fig. EV3A)

To prepare maximally phosphorylated MBP-XD2-C as a substrate for PP2A-B55, purified MBP-XD2-C (2 µM) was incubated with M-CDK (20 nM) and Suc1 (500 nM) in PR buffer at 22 °C for 60 min, followed by precipitation with anti-MBP beads (New England Biolabs, E8037). The bead-bound phosphorylated MBP-XD2-C (∼1 µM) was then incubated with PP2A-B55 (100 nM) in the presence or absence of okadaic acid (1 µM) at 22 °C for 60 min. Finally, aliquots of the reaction mixtures were analyzed by SDS-PAGE, followed by Pro-Q Diamond and CBB staining, or by immunoblotting to monitor the disappearance of the pT1353 phosphoepitope.

### Mitotic chromosome assembly and disassembly in *Xenopus* egg extracts

*Xenopus* egg extracts and demembranated sperm nuclei were prepared as described previously(Shintomi & Hirano, 2017, 2018). To recapitulate the events accompanying mitotic entry, sperm nuclei were added to interphase egg extracts at a final concentration of 1,000 nuclei/µl. After incubation at 22 °C for 60 min, the mixture was supplemented with non-degradable cyclin B1 (60 nM) to activate Cdk1 and further incubated for the indicated times. To mimic mitotic exit, sperm nuclei were first mixed with metaphase-arrested egg extracts, followed by the addition of CaCl₂ (500 µM) to trigger cyclin B degradation and consequent Cdk1 inactivation. At the indicated time points, aliquots were collected and analyzed by immunoblotting and immunofluorescence microscopy.

### Immunofluorescence microscopy and morphometric analysis

Reaction mixtures were fixed with 10 volumes of 4% formaldehyde in KMH fixative (20 mM Hepes-KOH [pH 7.7], 100 mM KCl, 2.5 mM MgCl_2_, 4% formaldehyde, 0.1% Triton X-100) and incubated at 22°C for 15 min. The mixtures were centrifuged at 3,000g for 10 min onto coverslips through a 15%-glycerol cushion containing KMH using a custom-made device to collect the fixed chromatin on a coverslip (Shintomi & Hirano, 2018). For double labeling using a combination of primary antibodies derived from mice and rabbits, the fixed chromatin on coverslips was incubated with a mixture of primary antibodies (2.0 µg/ml each) at 4°C overnight, followed by incubation with fluorescently labeled secondary antibodies (labeled with Alexa Fluor 488 or 568; 2.0 µg/ml) at room temperature for 2 h. After counterstaining with DAPI, the coverslips were mounted on glass slides with VectaShield mounting medium (H-1000, Vector Laboratories), sealed with nail polish, and processed for immunofluorescence microscopy.

Immunofluorescence microscopy was carried out using a DeltaVision microscope system (Cytiva), which consisted of an inverted fluorescence microscope (IX71, Olympus) equipped with a UPlanSApo 100×/1.40 oil-immersion lens and a CMOS camera (pco.edge 4.2m, PCO AG). Image stacks were acquired at a Z-step size of 0.2 µm and subjected to constrained iterative deconvolution with SoftWoRx software (version 2.0.0, Cytiva). Deconvolved image stacks were projected from three serial sections.

Quantitative morphometric analyses were performed using ImageJ/Fiji software (version 2.0.0-rc-43/1.52n, https://imagej.nih.gov/ij). Estimation of the DNA density was performed as follows: a chromatin-positive region was segmented in the DAPI channel using the Threshold function; the integrated density of DAPI within each segmented region and its area were measured with the Analyze Particles function; and DNA density was calculated by dividing the integrated density by the area. Other indices were estimated in a similar manner: for the individualization index, the perimeter of the region was divided by the integral DAPI intensity; for chromatin-bound condensin I and topo IIα, the integrated immunofluorescence intensity was normalized to that of DAPI; and for DNA standard deviation, the standard deviation of DAPI intensities in a segmented area was normalized to the mean DAPI value.

### AlphaFold structure prediction (Fig. EV4B)

AlphaFold3 multimer predictions were performed with the AlphaFold3 webserver (Abramson *et al*, 2024) using *X. laevis* CAP-G (residues 10-933) and CAP-H (residues 390-520, harboring a phosphorylated serine at position 415), and a 35-bp dsDNA fragment. Figures of structural models were generated with ChimeraX.

### Statistics and reproducibility

All data sets were processed using Excel (Excel for Mac version 16.x, Microsoft). Statistical analyses and graph drawing were carried out with Prism (version 9, GraphPad software). *P* values were assessed by an unpaired, two-tailed t-test with Welch’s correction. All experiments were independently reproduced at least three time, and representative results from these experiments are shown.

### Antibodies

An antibody specific to the phospho-epitope at S415 in XCAP-H (in-house identifier: AfR472-P) was raised against a synthetic peptide corresponding to the target region (EYSYFpSPRTMS; pS, phospho-serine). The following primary antibodies have been described previously (Hirano & Mitchison, 1994; Hirano *et al*, 1997; Kimura *et al*, 1998): anti-XCAP-C (AfR8), anti-XCAP-E (AfR9), anti-XCAP-D2 (AfR18), anti-XCAP-H (AfR20), and anti-pT1353-XCAP-D2 (AfR46-P).

Commercially available primary and secondary antibodies used in the current study were as follows: anti-topo II (MBL, M052-3 [RRID: AB_592894]); anti-nuclear core complex (Abcam, ab24609 [RRID: AB_448181]); Alexa Fluor 488-conjugated anti-mouse IgG (A11001 [RRID: AB_2534069], Thermo Fisher Scientific); Alexa Fluor 568-conjugated anti-rabbit IgG (A11036 [RRID: AB_10563566], Thermo Fisher Scientific); HRP-conjugated anti-rabbit IgG (PI-1000 [RRID: AB_2336198], Vector Laboratories).

## Acknowledgements

The authors thank Kazuhisa Kinoshita and Takao Ono for their discussion and comments. This study was supported by Grant-in-Aid for Scientific Research, KAKENHI (grants 25H02407 and 25K02211 [to K.S.]; 18H05276 and 20H05938 [to T.H.])

## Author contributions

**Keishi Shintomi:** Conceptualization; Data curation; Formal analysis; Funding acquisition; Investigation; Methodology; Project administration; Resources; Supervision; Validation; Visualization; Writing – original draft; Writing – review & editing. **Shoji Tane:** Conceptualization; Investigation; Methodology; Resources. **Yuki Masahara-Negishi:** Investigation; Methodology; Resources. **Tatsuya Hirao:** Conceptualization; Funding acquisition; Project administration; Supervision; Writing – original draft; Writing – review & editing.

## Disclosure and competing interest statement

The authors declare no competing interests.

**Figure EV1. Purification of full-length and tIDR-deleted versions of recombinant *X. laevis* condensin I complexes.**

**A.** Purification scheme of recombinant condensin I complexes. Aliquots were taken at the indicated steps and processed for the analyses shown in (B) and (C).

**B.** Upon purification of recombinant condensin I-FL and -ΔtIDRs, starting materials (fraction 1), intermediate fractions (2, 3, and 4), and final preparations (5) were analyzed by SDS-PAGE followed by Coomassie Brilliant Blue (CBB) staining.

**C.** 3C protease-treated eluates from the StrepTactin column were analyzed by SDS-PAGE before (fraction 3) and after (fraction 4) α-phosphatase treatment. The gel was first stained with Pro-Q Diamond to visualize phosphorylated proteins and then with CBB to visualize total proteins. Note that CAP-D2 and -H subunits were phosphorylated in host insect cells and that most, if not all, of these phosphate groups were removed by α-phosphatase.

**D.** Domain organization of *X. laevis* condensin I subunits. Cyclin-dependent kinase (Cdk) consensus sites (SP and TP motifs) identified in each subunit are indicated; sites conserved between *X. laevis* and humans are shown in red (18 out of 27 residues). Notably, some of these sites are clustered in the N-terminal IDR of CAP-H and the C-terminal IDR of CAP-D2. Because of the large molecular size of each subunit, comprehensive mapping of phosphorylation sites remains challenging. Nonetheless, phosphorylation of CAP-C, -D2, and CAP-H in *X. laevis* has been shown to depend on cyclin B-Cdk1 (Kimura *et al*, 1998; Shintomi *et al*, 2024). In particular, Cdk1 phosphorylation of T1314, T1348, and T1353 in CAP-D2 correlates with condensin I-mediated DNA supercoiling in vitro (Kimura *et al*, 1998). In human cells, alanine substitution of CAP-H S432 (corresponding to H-S415 in *X. laevis*) compromised condensin I binding to mitotic chromosomes (Kschonsak *et al*, 2017). Truncated versions of the non-SMC subunits lacking their terminal intrinsically disordered regions (tIDRs) are also shown.

**Figure EV2. Additional data on phosphorylation and chromatid reconstitution assays using recombinant condensin I complexes.**

**A-D.** Condensin I-FL_H-S415A and condensin I-ΔtIDRs_H-S415A were mock-incubated or incubated with M-CDK in the absence or presence of Suc1 in the indicated combinations. The reaction mixtures were analyzed as described in Fig. 1C-F.

**E.** An additional set of morphometric comparisons of chromatid reconstitution assays using condensin I-FL_H-S415A and -ΔtIDRs_H-S415A.

**F.** DNA relative standard deviation of chromatid reconstitution assays using condensin I-FL and condensin I-ΔtIDRs, as shown in Fig. 1. Notably, unlike the frost phenotype produced by condensin I-FL_H-S415A and condensin I-ΔtIDRs_H-S415A (Fig. 3C), chromatids produced by the complexes without the H-S415A mutations did not exhibit elevated DNA relative standard deviation.

Data information: In (B), data are presented as mean ± SEM from three independent experiments. In (E) and (F), data are presented as mean ± SD.

**Figure EV3. Inactivation of condensin I depends on the phosphatase activity of PP2A-B55.**

**A.** Validation of the phosphatase activity of purified PP2A-B55. A maltose-binding protein (MBP) fusion of an 80-residue fragment of *X. laevis* CAP-D2 (MBP-XD2-C), containing three TP motifs (T1314, T1348, and T1353), was phosphorylated with M-CDK in the presence of Suc1, and precipitated with anti-MBP beads. The bead-bound, maximally phosphorylated MBP-XD2-C was incubated with recombinant PP2A-B55 (100 nM) in the presence or absence of okadaic acid (1 µM). Reaction mixtures were analyzed by SDS-PAGE, followed by Pro-Q Diamond staining, CBB staining, and immunoblotting with peptide antibodies recognizing the XD2 C-tail and phosphorylated T1353.

**B.** After completion of mitotic chromatid assembly, reaction mixtures were incubated for an additional 100 min with PP2A-B55 (100 nM), either in the presence or absence of okadaic acid (1 µM). Chromatin was then fixed and subjected to immunofluorescence microscopy.

**Figure EV4. S415 in CAP-H is located near its DNA-binding motif.**

**A.** A model of condensin I in the ATP-free (apo) state bound to DNA, based on the cryo-EM structures of budding yeast condensin (Lee *et al*, 2022; Shaltiel *et al*, 2022). Condensin is proposed to bind to DNA through its two distinct modules, referred to as the motor and the anchor.

**B.** An AlphaFold3-predicted structure of the anchor-DNA complex, which is composed of the HEAT-repeat domain of *X. laevis* CAP-G (residues 10-933), the DNA-binding motif of *X. laevis* CAP-H known as the safety belt (residues 390-520, phosphorylated at S415)(Kschonsak *et al*, 2017), and a 35-bp dsDNA fragment. A close-up view (right) highlights the side chain of phosphorylated S415. The predicted structures showed no appreciable difference regardless of the phosphorylation status of H-S415.

## Notes

### Competing Interest Statement

The authors have declared no competing interest.

## References

Abramson J, Adler J, Dunger J, Evans R, Green T, Pritzel A, Ronneberger O, Willmore L, Ballard AJ, Bambrick J, et al (2024) Accurate structure prediction of biomolecular interactions with AlphaFold 3. Nature 630: 493–500

Chen J, Feng C, Wang Y & Chu X (2025) DNA actively regulates the “safety-belt” dynamics of condensin during loop extrusion. bioRxiv: 10.1101/2025.04.29.651239 [PREPRINT]

Chia KH, Takaki H, Fujimitsu K, Darling S, Zou J, Rappsilber J & Yamano H (2024) CDK1-PP2A-B55 interplay ensures cell cycle oscillation via Apc1-loop300. Cell Rep 43: 114155

Curran JF, Basu S, Auchynnikava T & Nurse P (2025) Control of CDK activity and the cell cycle by CKS proteins. bioRxiv: 10.1101/2025.07.04.663185

Cutts EE, Tetiker D, Kim E & Aragon L (2025) Molecular mechanism of condensin I activation by KIF4A. EMBO J 44: 682–704

Dekker B & Dekker J (2022) Regulation of the mitotic chromosome folding machines. Biochem J 479: 2153–2173

Eykelenboom JK, Gierliński M, Yue Z & Tanaka TU (2025) Nuclear exclusion of condensin I in prophase coordinates mitotic chromosome reorganization to ensure complete sister chromatid resolution. Curr Biol 35: 1562–1575.e7

Ganji M, Shaltiel IA, Bisht S, Kim E, Kalichava A, Haering CH & Dekker C (2018) Real-time imaging of DNA loop extrusion by condensin. Science 360: 102–105

Gibcus JH, Samejima K, Goloborodko A, Samejima I, Naumova N, Nuebler J, Kanemaki MT, Xie L, Paulson JR, Earnshaw WC, et al (2018) A pathway for mitotic chromosome formation. Science 359: eaaa06135

Godfrey M, Touati SA, Kataria M, Jones A, Snijders AP & Uhlmann F (2017) PP2ACdc55 phosphatase imposes ordered cell-cycle phosphorylation by opposing threonine phosphorylation. Mol Cell 65: 393–402.e3

Goloborodko A, Marko JF & Mirny LA (2016) Chromosome compaction by active loop extrusion. Biophys J 110: 2162–2168

Hirano T (2016) Condensin-based chromosome organization from bacteria to vertebrates. Cell 164: 847–857

Hirano T (2025) Mitotic genome folding. J Cell Biol 224: e202504075.

Hirano T, Kobayashi R & Hirano M (1997) Condensins, chromosome condensation protein complexes containing XCAP-C, XCAP-E and a Xenopus homolog of the Drosophila Barren protein. Cell 89: 511–521

Hirano T & Mitchison TJ (1994) A heterodimeric coiled-coil protein required for mitotic chromosome condensation in vitro. Cell 79: 449–458

Hoencamp C & Rowland BD (2023) Genome control by SMC complexes. Nat Rev Mol Cell Biol 24: 633–650

Kamenz J, Gelens L & Ferrell JE Jr (2021) Bistable, biphasic regulation of PP2A-B55 accounts for the dynamics of mitotic substrate phosphorylation. Curr Biol 31: 794–808.e6

Kimura K, Hirano M, Kobayashi R & Hirano T (1998) Phosphorylation and activation of 13S condensin by Cdc2 in vitro. Science 282: 487–490

Kschonsak M, Merkel F, Bisht S, Metz J, Rybin V, Hassler M & Haering CH (2017) Structural basis for a safety-belt mechanism that anchors condensin to chromosomes. Cell 171: 588–600.e24

Lee B-G, Merkel F, Allegretti M, Hassler M, Cawood C, Lecomte L, O’Reilly FJ, Sinn LR, Gutierrez-Escribano P, Kschonsak M, et al (2020) Cryo-EM structures of holo condensin reveal a subunit flip-flop mechanism. Nat Struct Mol Biol 27: 743–751

Lee B-G, Rhodes J & Löwe J (2022) Clamping of DNA shuts the condensin neck gate. Proc Natl Acad Sci U S A 119: e2120006119

Magalska A, Schellhaus AK, Moreno-Andrés D, Zanini F, Schooley A, Sachdev R, Schwarz H, Madlung J & Antonin W (2014) RuvB-like ATPases function in chromatin decondensation at the end of mitosis. Dev Cell 31: 305–318

McGrath DA, Balog ERM, Kõivomägi M, Lucena R, Mai MV, Hirschi A, Kellogg DR, Loog M & Rubin SM (2013) Cks confers specificity to phosphorylation-dependent CDK signaling pathways. Nat Struct Mol Biol 20: 1407–1414

Mochida S, Ikeo S, Gannon J & Hunt T (2009) Regulated activity of PP2A-B55 delta is crucial for controlling entry into and exit from mitosis in Xenopus egg extracts. EMBO J 28: 2777–2785

Murray AW (1991) Cell cycle extracts. Methods Cell Biol 36: 581–605

Nash P, Tang X, Orlicky S, Chen Q, Gertler FB, Mendenhall MD, Sicheri F, Pawson T & Tyers M (2001) Multisite phosphorylation of a CDK inhibitor sets a threshold for the onset of DNA replication. Nature 414: 514–521

Ono T, Losada A, Hirano M, Myers MP, Neuwald AF & Hirano T (2003) Differential contributions of condensin I and condensin II to mitotic chromosome architecture in vertebrate cells. Cell 115: 109–121

Paulson JR, Hudson DF, Cisneros-Soberanis F & Earnshaw WC (2021) Mitotic chromosomes. Semin Cell Dev Biol 117: 7–29

Rata S, Suarez Peredo Rodriguez MF, Joseph S, Peter N, Echegaray Iturra F, Yang F, Madzvamuse A, Ruppert JG, Samejima K, Platani M, et al (2018) Two interlinked bistable switches govern mitotic control in mammalian cells. Curr Biol 28: 3824–3832.e6

Robellet X, Thattikota Y, Wang F, Wee T-L, Pascariu M, Shankar S, Bonneil É, Brown CM & D’Amours D (2015) A high-sensitivity phospho-switch triggered by Cdk1 governs chromosome morphogenesis during cell division. Genes Dev 29: 426–439

Saitoh N, Goldberg IG, Wood ER & Earnshaw WC (1994) ScII: an abundant chromosome scaffold protein is a member of a family of putative ATPases with an unusual predicted tertiary structure. J Cell Biol 127: 303–318

Shaltiel IA, Datta S, Lecomte L, Hassler M, Kschonsak M, Bravo S, Stober C, Ormanns J, Eustermann S & Haering CH (2022) A hold-and-feed mechanism drives directional DNA loop extrusion by condensin. Science 376: 1087–1094

Shintomi K & Hirano T (2011) The relative ratio of condensin I to II determines chromosome shapes. Genes Dev 25: 1464–1469

Shintomi K & Hirano T (2017) A sister chromatid cohesion assay using Xenopus egg extracts. Methods Mol Biol 1515: 3–21

Shintomi K & Hirano T (2018) Reconstitution of mitotic chromatids in vitro. Curr Protoc Cell Biol 79: e48

Shintomi K & Hirano T (2021) Guiding functions of the C-terminal domain of topoisomerase IIα advance mitotic chromosome assembly. Nat Commun 12: 2917

Shintomi K, Masahara-Negishi Y, Shima M, Tane S & Hirano T (2024) Recombinant cyclin B-Cdk1-Suc1 capable of multi-site mitotic phosphorylation in vitro. PLoS One 19: e0299003

Shintomi K, Takahashi TS & Hirano T (2015) Reconstitution of mitotic chromatids with a minimum set of purified factors. Nat Cell Biol 17: 1014–1023

St-Pierre J, Douziech M, Bazile F, Pascariu M, Bonneil E, Sauvé V, Ratsima H & D’Amours D (2009) Polo kinase regulates mitotic chromosome condensation by hyperactivation of condensin DNA supercoiling activity. Mol Cell 34: 416–426

Sutani T, Yuasa T, Tomonaga T, Dohmae N, Takio K & Yanagida M (1999) Fission yeast condensin complex: essential roles of non-SMC subunits for condensation and Cdc2 phosphorylation of Cut3/SMC4. Genes Dev 13: 2271–2283

Takemoto A, Kimura K, Yokoyama S & Hanaoka F (2004) Cell cycle-dependent phosphorylation, nuclear localization, and activation of human condensin. J Biol Chem 279: 4551–4559

Tane S, Shintomi K, Kinoshita K, Tsubota Y, Yoshida MM, Nishiyama T & Hirano T (2022) Cell cycle-specific loading of condensin I is regulated by the N-terminal tail of its kleisin subunit. Elife 11: e84694

Tang M, Pobegalov G, Tanizawa H, Chen ZA, Rappsilber J, Molodtsov M, Noma K-I & Uhlmann F (2023) Establishment of dsDNA-dsDNA interactions by the condensin complex. Mol Cell 83: 3787–3800.e9

Thadani R, Kamenz J, Heeger S, Muñoz S & Uhlmann F (2018) Cell-cycle regulation of dynamic chromosome association of the condensin complex. Cell Rep 23: 2308–2317

Thattikota Y, Tollis S, Palou R, Vinet J, Tyers M & D’Amours D (2018) Cdc48/VCP promotes chromosome morphogenesis by releasing condensin from self-entrapment in chromatin. Mol Cell 69: 664–676.e5

Tsubota Y, Shintomi K, Kinoshita K, Masahara-Negishi Y, Aizawa Y, Shima M, Hirano T & Nishiyama T (2025) Functional interplay between condensin I and topoisomerase IIα in single-molecule DNA compaction. Nat Commun 16: 7239

Vagnarelli P (2021) Back to the new beginning: Mitotic exit in space and time. Semin Cell Dev Biol 117: 140–148

Vagnarelli P, Hudson DF, Ribeiro SA, Trinkle-Mulcahy L, Spence JM, Lai F, Farr CJ, Lamond AI & Earnshaw WC (2006) Condensin and Repo-Man-PP1 co-operate in the regulation of chromosome architecture during mitosis. Nat Cell Biol 8: 1133–1142

Yeong FM, Hombauer H, Wendt KS, Hirota T, Mudrak I, Mechtler K, Loregger T, Marchler-Bauer A, Tanaka K, Peters J-M, et al (2003) Identification of a subunit of a novel Kleisin-beta/SMC complex as a potential substrate of protein phosphatase 2A. Curr Biol 13: 2058–2064

Yoshida MM, Kinoshita K, Aizawa Y, Tane S, Yamashita D, Shintomi K & Hirano T (2022) Molecular dissection of condensin II-mediated chromosome assembly using in vitro assays. Elife 11: e78984

Yoshida MM, Kinoshita K, Shintomi K, Aizawa Y & Hirano T (2024) Regulation of condensin II by self-suppression and release mechanisms. Mol Biol Cell 35: ar21

Zeisner TU, Auchynnikava T & Nurse P (2025) Phosphatase specificity influences phosphorylation timing of CDK substrates during the cell cycle. bioRxiv: 10.1101/2025.02.21.639482 [PREPRINT]

