## Supplementary figures and images for "Reconstitution of phospho-regulated mitotic chromatid assembly and disassembly"

### Supplemental Figure EV1

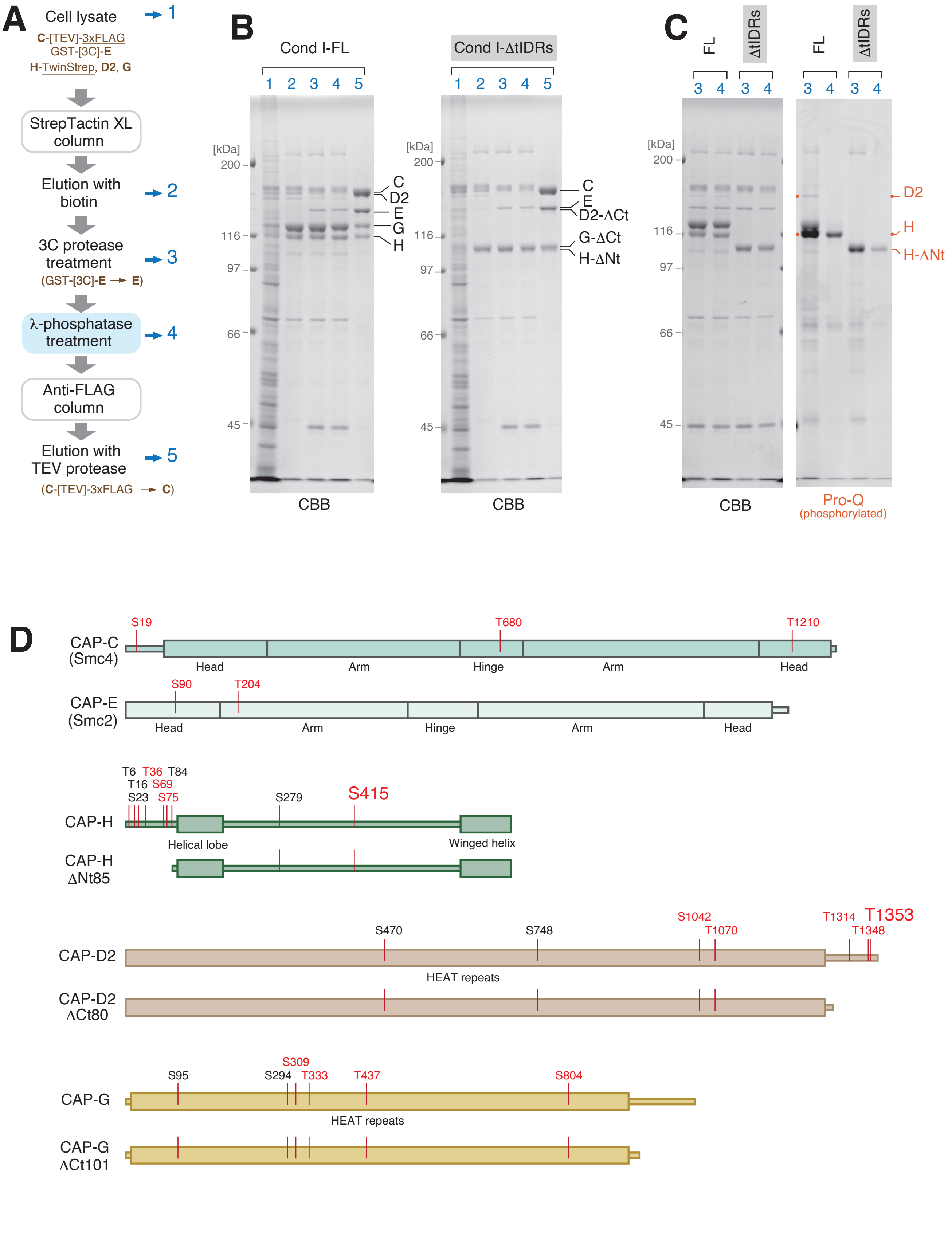


**Fig. EV1_Shintomi et al**

### Supplemental Figure EV2

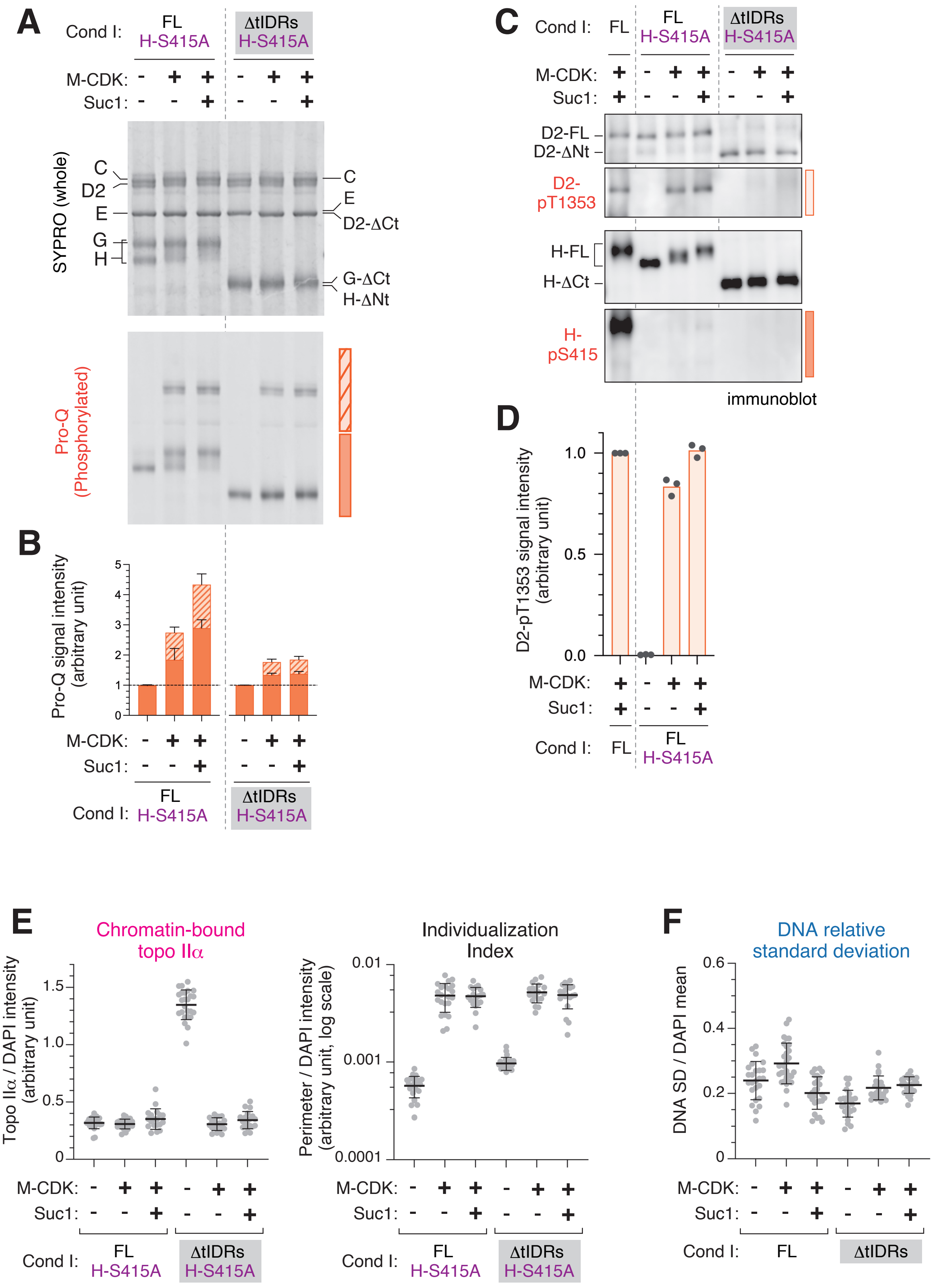


#### Fig. EV2_Shintomi et al

### Supplemental Figure EV3

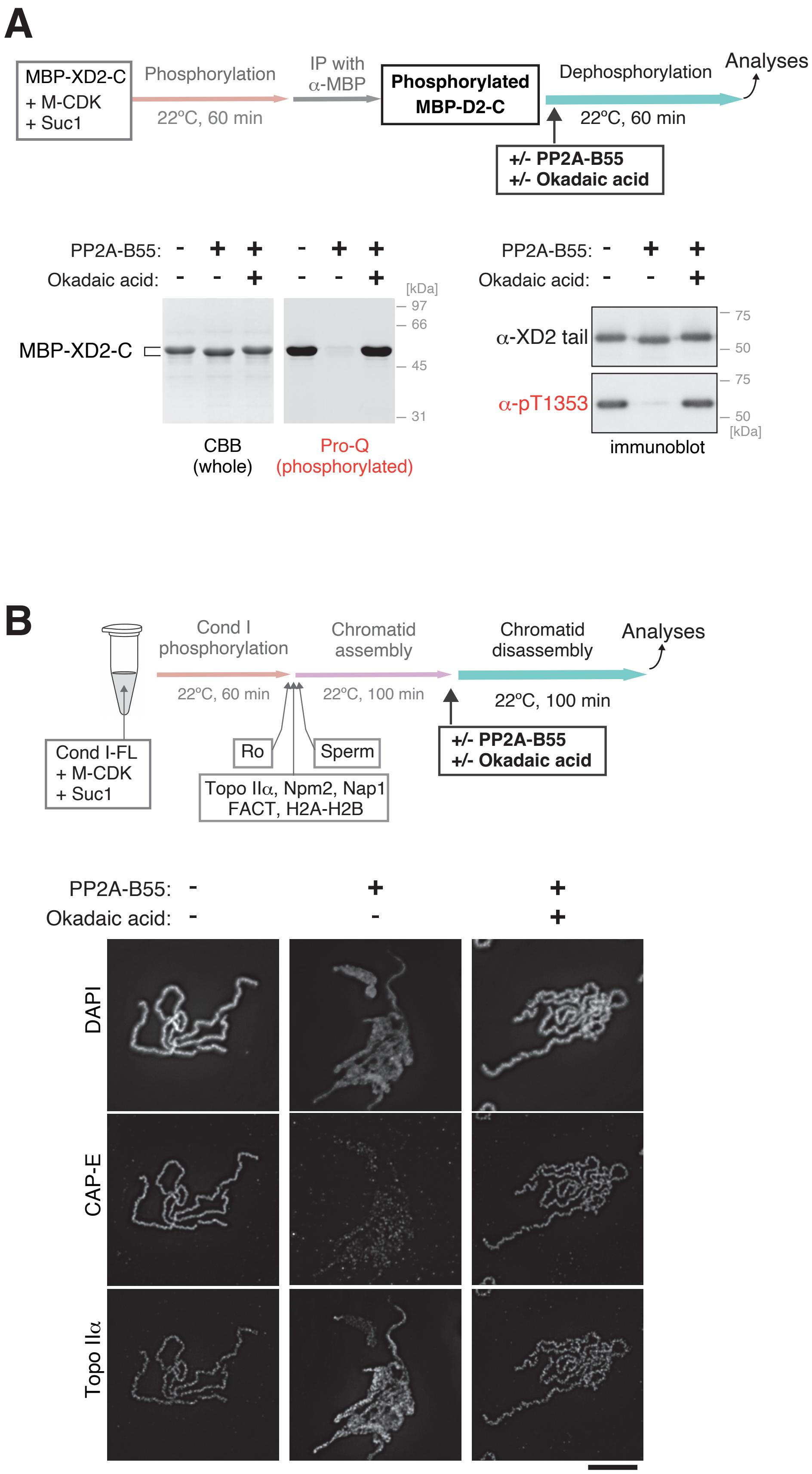


#### Fig. EV3_Shintomi et al

### Supplemental Figure EV4

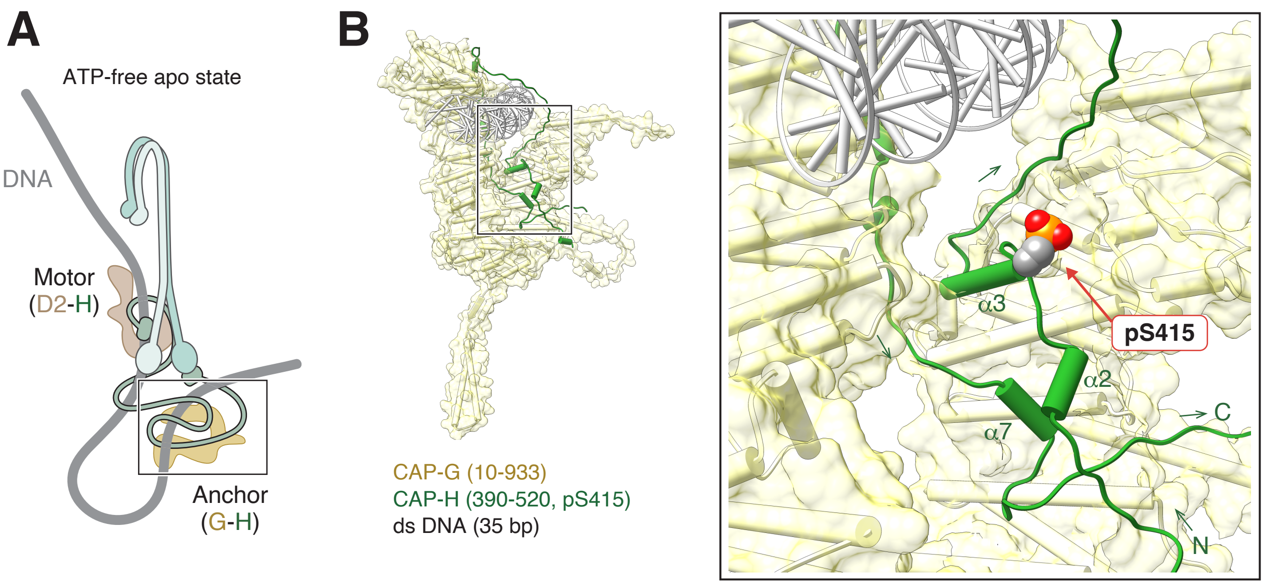


#### Fig. EV4_Shintomi et al
